# Endogenous oxytocin-linked heart rate synchrony for social connection in live sports spectators

**DOI:** 10.1101/2025.09.15.676321

**Authors:** Takashi Matsui, Takafumi Yamaguchi, Daisuke Funabashi, Shion Takahashi, Hiroki Matsuoka, Shohei Dobashi, Kiyonobu Kigoshi, Shinzo Yamada, Hideki Takagi

## Abstract

Oxytocin has been proposed as a pharmacological target for social dysfunction. However, clinical trials of exogenous oxytocin have yielded modest and inconsistent benefits, highlighting the need for naturalistic, non-pharmacological approaches that recruit endogenous oxytocin. This prospective observational field study examined whether live sports spectating in a modest collegiate setting engages endogenous oxytocin and heart rate synchrony, and how these processes relate to social bonding. Sixty casual spectators at two university basketball games provided salivary samples for oxytocin and cortisol and completed psychological questionnaires before the game, at halftime, and after the game, while heart rate was continuously recorded. Oxytocin dynamics were baseline dependent: individuals with low pre-game oxytocin showed clear increases during spectating, whereas those with high pre-game oxytocin maintained elevated concentrations; across both groups, cortisol decreased and the oxytocin-to-cortisol ratio increased, indicating a shift toward an oxytocin-dominant, low-stress hormonal profile. Live spectating also enhanced heart rate synchrony between spectators, and higher oxytocin levels and stronger synchrony were associated with greater enjoyment, stronger unity with players and fellow fans, higher flow, and stronger intention to revisit. Mediation analyses indicated that perceived unity mediated links between physiological responses and experiential outcomes. These findings identify coordinated oxytocin dynamics and physiological synchrony as candidate mechanisms through which live sports spectating can foster social connection and positive psychological states in everyday, low-stakes settings. Live sports events may represent a simple, scalable non-pharmacological approach to engaging oxytocin-related systems and interpersonal synchrony to promote social connection in everyday life.

## 1. Introduction

Oxytocin is a hypothalamic neuropeptide with potent central effects on social cognition and stress regulation, and has attracted considerable interest as a pharmacological target for social dysfunction in neuropsychiatric disorders such as autism spectrum disorder and schizophrenia[1–3]. Exogenous oxytocin administered intranasally can acutely modulate social attention, emotion recognition, and activity in limbic–striatal circuits, and has therefore been trialled as an adjunctive treatment for social impairments [4–6]. However, large-scale and longer-term clinical trials have yielded small or inconsistent benefits on core social symptoms, and considerable uncertainty remains regarding optimal dosing regimens, individual response profiles, and long-term safety [7–9]. In addition, peripherally administered oxytocin has an unfavourable pharmacokinetic and delivery profile, including a short plasma half-life, limited central penetration, and off-target peripheral effects, and even though its short-term safety profile appears acceptable, rare adverse reactions have been reported with repeated or high-dose use [10,11]. These limitations have stimulated growing interest in naturalistic, non-pharmacological strategies that can recruit endogenous oxytocin to engage oxytocin-linked circuits for social connection without the constraints of exogenous drug administration [12,13].

Beyond its classical roles in parturition and lactation, endogenous oxytocin is released by everyday affiliative cues, particularly low-intensity, non-noxous social touch such as gentle stroking or massage, which has been linked to stress-buffering effects and increased well-being [14,15]. Experimental studies in humans show that affective tactile stimulation, including massage-like foot stimulation, can acutely increase peripheral oxytocin concentrations and concomitantly engage cortical regions implicated in social cognition and reward, such as the orbitofrontal cortex and superior temporal sulcus [14,16]. Similarly, traditional martial arts training, which combines vigorous physical exercise with structured dyadic contact, increases salivary oxytocin immediately after high-intensity sparring, with larger oxytocin responses during close-contact ground grappling than during more distal “punch–kick” sparring [17]. Martial-arts–based interventions in youths at high psychosocial risk further indicate that oxytocin and cortisol responses during training differ between high-risk and low-risk groups, with high-risk youths showing lower baseline oxytocin levels and altered reactivity compared with low-risk peers [18].

Familiar vocal social contact can also trigger oxytocin release after stress; for example, comfort delivered solely via a mother’s voice elevates oxytocin and accelerates cortisol recovery in children, suggesting that social vocalizations provide a potent non-tactile route for engaging oxytocin-linked bonding and stress-regulation pathways [19,20]. Endogenous oxytocin levels are likewise elevated in naturally occurring affiliative contexts such as the early stages of romantic attachment, where higher plasma oxytocin covaries with couples’ interactive reciprocity and dyadic synchrony [21]. A recent systematic review and meta-analysis concluded that endogenous oxytocin concentrations show a modest but reliable association with human social interaction overall, while also revealing substantial heterogeneity across individuals, interaction types, and assay methods [22,23]. Collectively, these findings suggest that multiple sensory, motor, and relational channels, including social touch, vocal communication, and structured physical practices, can recruit endogenous oxytocin in everyday life, but they also underscore the need to identify robust, ecologically valid social behaviours that reliably engage the human oxytocin system despite this heterogeneity and methodological uncertainty [22].

Large-scale collective events such as religious rituals, concerts, and sporting matches can generate intense shared emotional experiences and feelings of togetherness, and recent work suggests that physiological synchrony in crowds may be one mechanism through which such events promote social bonding [24,25]. Field and laboratory studies show that being part of a crowd or dyad is associated with increased heart rate synchrony, and that greater physiological synchrony predicts stronger feelings of connectedness and cooperative attitudes [26,27]. Such increases in synchrony are likely driven not only by individual arousal but also by shared external stimuli, including the tension of the game, the roar of the crowd, and collective emotional reactions, that act as common drivers of cardiac dynamics across individuals. In parallel, social psychological research on sport fandom has developed the Team Identification–Social Psychological Health Model, which posits that identifying with a team and engaging in spectatorship help fans gain social connections that, in turn, enhance social well-being [28]

Consistent with this framework, observational and longitudinal studies indicate that live sport spectatorship and stronger team loyalty are associated with greater life satisfaction, perceived emotional support from other fans, and reduced loneliness, particularly among middle-aged and older adults [29,30]. From a neuroendocrine perspective, theoretical work has proposed that oxytocin may support coordination, trust, and cohesion in team sport environments, yet empirical research in sports has focused predominantly on cortisol and testosterone responses to competition in athletes and, to a lesser extent, fans [31,32]. Despite growing interest in the psychobiology of fandom, oxytocin in spectators remains largely unexplored. To date, however, no study has simultaneously examined endogenous oxytocin dynamics, cardiac synchrony among co-present spectators, and changes in perceived social bonding within a real-world live sports event, particularly among casual fans who do not belong to established supporter groups.

We therefore hypothesized that a single session of live sports spectating by casual spectators induces baseline-dependent increases in endogenous oxytocin in association with enhancements in interpersonal heart rate synchrony, thereby promoting post-game unity with fellow fans and loyalty to the team. To test this hypothesis, we conducted a field study of previously unacquainted casual spectators at collegiate basketball games, integrating repeated measures of salivary oxytocin and cortisol with continuous heart rate recording and self-reported indices of social connection and experiential quality.

## 2. Materials and Methods

### 2.1. Participants

We conducted a prospective observational field study with spectators attending home basketball games at the University of Tsukuba. A total of 62 adults were recruited from attendees of “TSUKUBA LIVE!”, the university’s home event series (Fig. 1A). Unlike professional sports events, these collegiate games attracted modest crowds primarily composed of students and local residents, reflecting a predominantly casual rather than highly devoted fan base. Each participant attended one game, which resulted in either a win or a loss for the home team.

**Figure 1.**
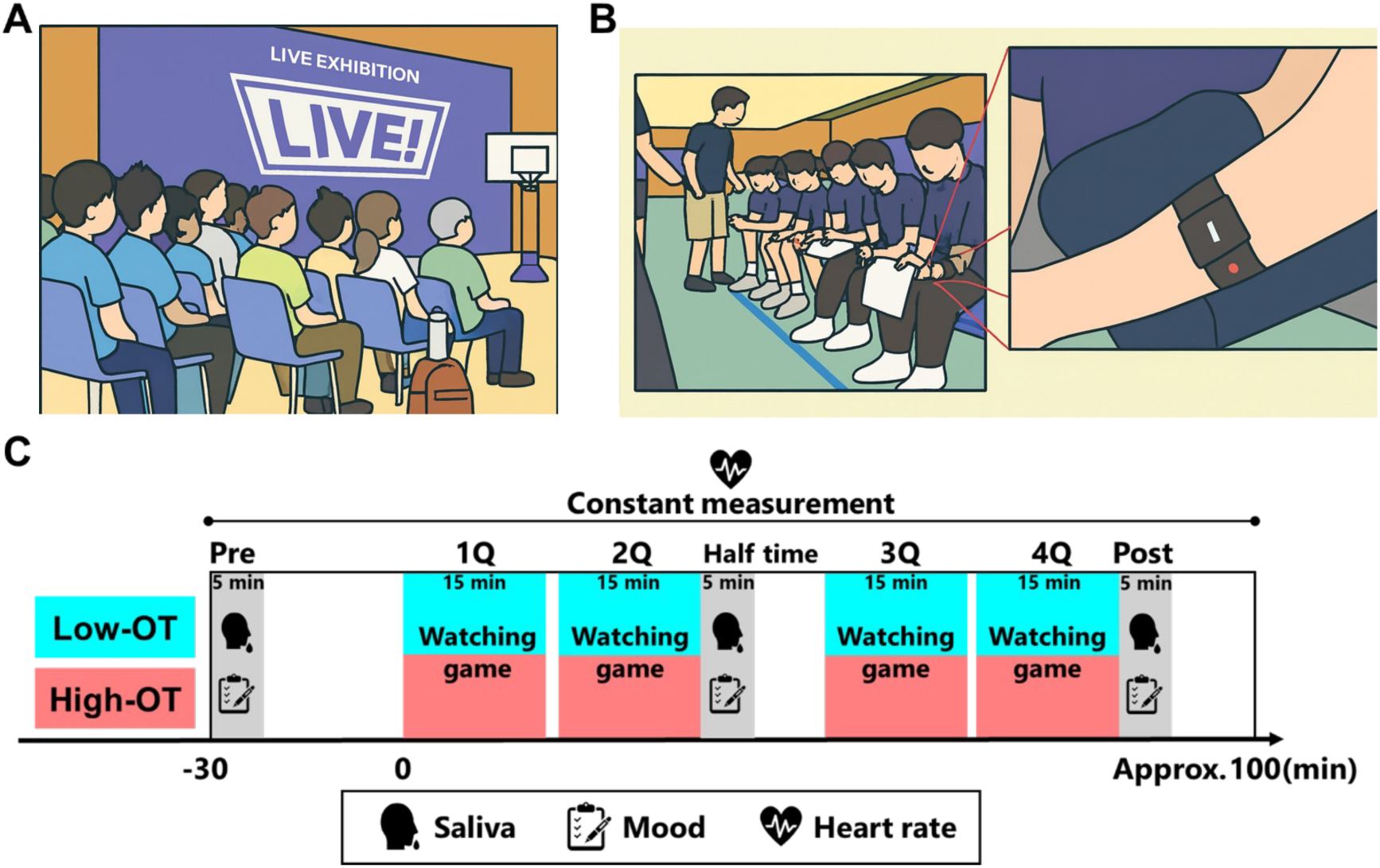
The experimental protocol. **A.** A typical photo of the spectating state. **B.** A typical photo of saliva sampling, questionnaire completion, and the HR measurement device. **C.** Overview of the experimental protocol. The participants watch basketball games for approximately 100 minutes. Saliva samples and mood questionnaires were administered before. Each measurement took approximately 5 minutes to complete. The HR was measured during spectating.

Two participants were excluded from all analyses due to insufficient pre-game saliva samples, leaving 60 participants in the final analytic sample. Based on the median pre-game salivary oxytocin concentration (27.51 pg/mL; Supplementary Table 1), participants were classified into a high baseline oxytocin group (High-OT; n = 30; 22.2 ± 8.6 years; 10 females) and a low baseline oxytocin group (Low-OT; n = 30; 22.9 ± 9.0 years; 11 females). Baseline oxytocin levels were higher in the High-OT group than in the Low-OT group by design, whereas there were no significant group differences in sex, age, body composition, resting heart rate, or habitual physical activity, indicating that the grouping primarily reflected pre-game oxytocin concentration rather than broader demographic or physiological differences (Table S1).

An a priori power analysis was conducted using G*Power 3.1, based on a partial η² of 0.036 derived from a previous study examining the association between baseline oxytocin levels and responses to video viewing [33]. This analysis indicated that a total sample size of 60 participants would provide 90% power to detect a significant difference between two independent means in a two-sided test with an α level of 0.05.

Inclusion criteria were as follows: age 18 years or older; ability to understand the study procedures and provide informed consent; no current medication for neuropsychiatric, cardiovascular, or metabolic diseases; and ability to attend the stadium for the duration of the measurements. The study was approved by the Research Ethics Committee of the Institute of Health and Sport Sciences, University of Tsukuba, and was conducted in accordance with the Declaration of Helsinki. All participants provided written informed consent prior to participation. The study is reported in accordance with the STROBE guidelines for observational studies.

### 2.2. Study design and procedure

This prospective observational field study was conducted during two home basketball games at the University of Tsukuba (Fig. 1C). The study was designed to capture coupled endocrine and cardiovascular responses to live sports spectating by combining repeated saliva sampling with continuous heart rate recording and self-report measures of mood and social connection.

When inviting participants, an outline of the study and a URL link to an online pre-survey (see Section 2.3) were sent via e-mail to potential spectators of the home event series “TSUKUBA LIVE!”. Participants were instructed to refrain from consuming alcohol from the day before the game until the end of the experiment, to avoid caffeine intake and strenuous physical exercise on the day of the game, and to abstain from eating for at least 2 hours before arriving at the stadium.

On the day of measurement, participants arrived at the indoor arena approximately 60 minutes before tipoff. Upon arrival, they received a detailed explanation of the procedures again and provided written informed consent. Participants were then fitted with an optical heart rate sensor (Polar Verity Sense; Polar, Finland) on the upper arm and completed baseline saliva sampling (2 mL) for hormone analysis as well as baseline psychological questionnaires in the concourse area (Fig. 1B). After these baseline assessments, participants moved to their assigned seats and watched the game under naturalistic conditions. Additional saliva samples and questionnaires were collected at halftime and immediately after the game, yielding three repeated measurement points: pre-game (T1), halftime (T2), and post-game (T3).

### 2.3. Demographic and physical activity assessment

Before the game, participants completed an online pre-survey delivered via Google Forms to collect demographic characteristics (for example, age and sex) and habitual physical activity. Habitual physical activity was assessed using the Japanese version of the International Physical Activity Questionnaire (IPAQ), which estimates average daily time spent in vigorous-intensity, moderate-intensity, and walking activities, as well as sedentary behaviour.

Some participants did not complete all demographic or physical activity items, resulting in missing data for certain questionnaire variables. These missing values were not imputed. Analyses involving those specific variables were conducted using available-case data, whereas participants’ physiological (heart rate, salivary hormones) and psychological (questionnaire) measures were retained for all other analyses when data quality met the inclusion criteria.

### 2.4. Physiological measurements and saliva collection

To evaluate the physiological impact of live sports spectating, we assessed cardiovascular activity and salivary hormone concentrations (oxytocin and cortisol) across the three measurement points (pre-game, halftime, post-game). Heart rate (HR) was monitored continuously using a standalone optical HR sensor worn on the upper arm (Polar Verity Sense; Polar, Finland), as described in Section 2.2. Participants wore the sensor from approximately 30 minutes before tipoff until 15 minutes after the end of the game. HR data were sampled at 1 Hz throughout this period. For subsequent analyses, the spectating period was segmented into “in-game” intervals (when play was in progress and players were on the court) and “out-of-game” intervals (inter-quarter breaks, halftime, and the 15-minute post-game period).

Salivary oxytocin and cortisol were measured as neuroendocrine markers indexing social bonding and stress-related responses during spectating. At each time point (pre-game, halftime, post-game), participants provided approximately 2 mL of whole saliva by passively drooling through a straw directly into polypropylene collection tubes. Samples were immediately placed on ice and then stored at −80 °C until processing. To precipitate mucins and other debris, frozen samples were thawed and centrifuged at 1500 × g for 20 minutes; the clear supernatant was then aliquoted and re-frozen at −80 °C until assay.

For oxytocin quantification, 1000 μL of each saliva sample was concentrated by centrifugal evaporation and reconstituted according to the manufacturer’s instructions before assay with a commercial competitive enzyme-linked immunosorbent assay (ELISA) kit (Oxytocin ELISA Kit, Enzo Life Sciences, USA). Salivary cortisol concentration was measured using a high-sensitivity ELISA kit (Salivary Cortisol ELISA Kit, Salimetrics, USA). All samples were assayed in duplicate on 96-well plates, and absorbance was read using a multimode microplate reader (Varioskan LUX, Thermo Fisher Scientific, USA). Hormone concentrations were calculated from standard curves generated for each plate. Across all assays, the coefficient of variation for duplicate determinations was below 5%, indicating acceptable analytic reliability.

### 2.5. HR data analysis

Heart rate synchrony between spectators was quantified using cross-correlation function (CCF) analysis of the HR time series [34]. For each participant pair, CCFs were computed within a temporal lag window of ±30 seconds to identify the maximum correlation between their HR time series. The temporal lag at which the maximum CCF occurred was extracted, and the absolute value of this lag was taken as an index of the degree of temporal alignment irrespective of direction, with smaller values indicating tighter alignment.

Because participants were seated in two separate blocks on opposite sides of the same courtside, their visual perspectives of the game could differ and potentially influence synchrony. To account for this, synchrony scores were calculated within each seating block rather than across all participants. For each participant, an individual synchrony score was obtained by averaging the dyadic CCF values across all pairings with other participants in the same seating block.

Baseline HR synchrony was defined as the 5-minute interval during the pre-game period that showed the lowest within-block CCF values, and synchrony indices during spectating were expressed relative to this baseline where appropriate. In-game and out-of-game synchrony measures were derived from the segmented HR data described in Section 2.4, enabling us to examine whether physiological synchrony differed between active play and non-play intervals and how these dynamics related to neuroendocrine and psychological measures.

### 2.6. Psychological questionnaires

Subjective experiences during spectating were assessed using four self-report measures: enjoyment, flow state, intention to revisit, and perceived social unity.

**Enjoyment.** Momentary enjoyment while watching the game was assessed using a 100-mm visual analogue scale (VAS). The anchors “not enjoyable at all” and “very enjoyable” were printed at the left and right ends, respectively. Participants were instructed to place an “X” on the line at the point that best represented how much they were enjoying the game at that moment. The distance (mm) from the left anchor to the mark was measured and expressed as a percentage of the total line length, with higher scores indicating greater enjoyment.

**Flow state.** Flow experience during spectating was measured using a brief four-item flow scale developed for this study, which asked participants the extent to which they (1) forgot the time, (2) were completely focused, (3) lost themselves, and (4) enjoyed themselves while watching the game. Each item was rated on a seven-point Likert scale ranging from 1 (“not at all”) to 7 (“very much”), and the sum of the four items was used as an index of the subjective flow state during spectating.

**Intention to revisit.** To capture behavioural intention to re-engage with live sports spectating, we administered a single-item loyalty scale that assessed intention to revisit: “I would like to watch this event again in the future.” Responses were given on a seven-point Likert scale from 1 (“not at all”) to 7 (“very much”), with higher scores indicating stronger intention to revisit.

**Sense of unity (Inclusion of Other in the Self).** Perceived social unity with players and fellow spectators was assessed using the Inclusion of Other in the Self (IOS) scale, a widely used pictorial measure of interpersonal closeness [35]. The IOS presents seven pairs of overlapping circles; greater overlap indicates a closer perceived relationship. In this study, participants rated their perceived closeness to four targets: home team players, home team spectators, away team players, and away team spectators. For each target, participants selected the pair of circles that best represented their current sense of unity with that group.

Enjoyment (VAS) and unity (IOS) were assessed at all three time points (pre-game, halftime, post-game) to capture temporal changes across spectating. Flow state and intention to revisit were measured only immediately after the game, reflecting participants’ overall experiential and behavioural responses to the event.

### 2.7 Statistical analysis

Heart rate synchrony analyses and mediation models were conducted using R statistical software (version 4.4.0; R Foundation for Statistical Computing, Vienna, Austria). All other statistical analyses were performed using GraphPad Prism version 10 (GraphPad Software, San Diego, CA, USA).

Depending on the data structure, group differences and temporal changes were examined using unpaired t-tests or two-way analyses of variance (ANOVA) with factors for group (High-OT vs. Low-OT) and time (pre-game, halftime, post-game). When Mauchly’s test indicated violation of sphericity, Greenhouse–Geisser corrections were applied to the degrees of freedom. When a significant main effect or interaction was detected, Bonferroni’s multiple comparison test was used for post hoc comparisons.

In cases where missing values arose due to incomplete questionnaire responses or exclusion of physiologically implausible outliers, linear mixed-effects models were employed instead of repeated-measures ANOVA to maximise the use of available data. For mixed-effects models, F-statistics and degrees of freedom were used to derive partial η² as an approximate index of effect size. Correlation analyses, for example associations between oxytocin, heart rate synchrony indices, and psychological measures, were conducted using Pearson’s product–moment correlation coefficient unless otherwise specified.

For mediation analyses, indirect effects were estimated using a nonparametric bootstrapping procedure with 2000 resamples and bias-corrected 95 per cent confidence intervals, implemented in the R package lavaan. An indirect effect was considered statistically significant when the 95 per cent confidence interval did not include zero. All data are presented as mean ± standard deviation (SD) unless stated otherwise, and the threshold for statistical significance was set at p < 0.05 (two-tailed).

## 3. Results

### 3.1. Endogenous oxytocin dynamics shift in a baseline-dependent manner during live sports spectating

We first tested whether live sports spectating elicits distinct hormonal responses depending on baseline oxytocin levels. The High-OT group maintained elevated oxytocin concentrations throughout the game, showing no evidence of further time-dependent change, whereas the Low-OT group exhibited a significant increase over time that brought post-game oxytocin levels into the range observed in the High-OT group (High-OT main group effect: F₁,₅₈ = 29.06, p < 0.0001, ηp² = 0.33; Low-OT time effect: F₂,₁₁₆ = 11.67, p < 0.0001, ηp² = 0.16; Fig. 2A). Cortisol concentrations decreased across both groups during spectating (F₁.₈,₁₀₂.₇ = 3.32, p = 0.045, ηp² = 0.05; Fig. 2B), resulting in a progressive increase in the oxytocin-to-cortisol ratio (F₁.₇,₉₅.₇ = 14.50, p < 0.0001, ηp² = 0.20; Fig. 2C). Taken together, these findings indicate that sports spectating modulates hormonal dynamics in a baseline-dependent manner, sustaining high oxytocin levels in one subgroup and augmenting them in another, while shifting both groups toward an oxytocin-dominant, low-stress hormonal balance without evidence of stress-related endocrine activation.

**Figure 2.**
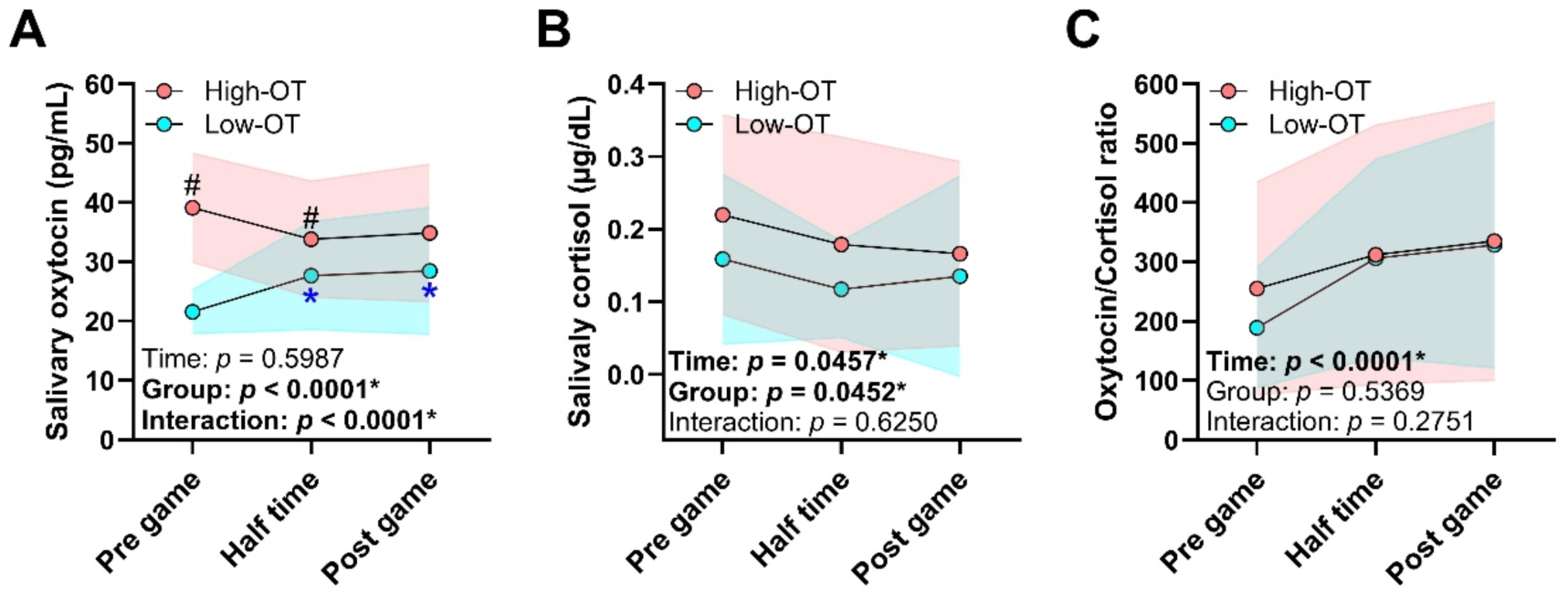
Endogenous oxytocin dynamics shift in a baseline-dependent manner during live sports spectating. **A.** saliva oxytocin levels. **B.** saliva cortisol levels. **C.** oxytocin/cortisol ratio. Standard divisions are shown as red (High-OT) bands and blue (Low-OT) on the graph. Results of two-way ANOVA are shown in the lower left of each graph. Time: main effect of watching time. Group: main effect of the group. Interaction: interaction between time and group. *: p < 0.05 vs. pre game for both groups; #: p < 0.05 vs. Low-OT.

### 3.2. High oxytocin promotes unity through enhanced enjoyment

We next examined how oxytocin dynamics during spectating relate to subjective experiences, focusing on social bonding and enjoyment. Unity with the home team players and home team fans increased over time irrespective of baseline oxytocin group (players: F₁.₄₈,₈₄.₄₁ = 24.39, p < 0.0001, ηp² = 0.30; fans: F₁.₈₂,₁₀₄.₁₀ = 12.36, p < 0.0001, ηp² = 0.17; Fig. 3A, B). Enjoyment also increased in both groups (F₁.₇₉,₁₀₁.₇₀ = 22.18, p < 0.0001, ηp² = 0.28), with a more pronounced effect in the High-OT group than in the Low-OT group (group effect: F₁,₅₇ = 6.59, p = 0.012, ηp² = 0.10; Fig. 3C).

**Figure 3.**
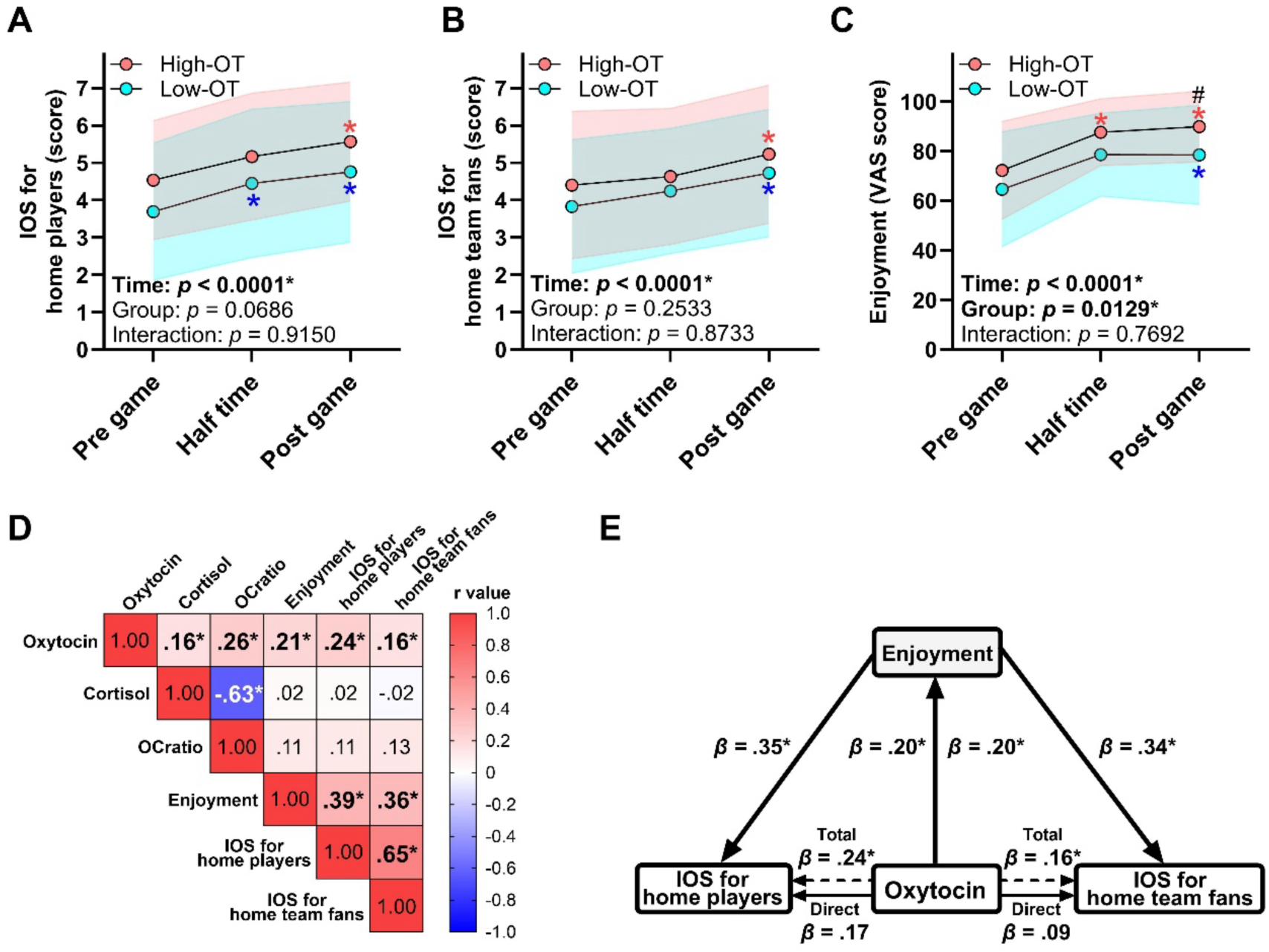
High oxytocin promotes unity through enhanced enjoyment. **A, B.** A sense of unity with home players (A) and home team fans (B). **C**. Enjoyment. Standard divisions are shown as red (High-OT) bands and blue (Low-OT) on the graph. Results of two-way ANOVA are shown in the lower left of each graph. Time: main effect of watching time. Group: main effect of the group. Interaction: interaction between time and group. *: p < 0.05 vs. pre game for both groups; #: p < 0.05 vs. Low-OT. **D**. Correlation matrix summarizing associations between neuroendocrine responses, mood and a sense of unity. **E**. Mediation model showing the associations between oxytocin, enjoyment, and a sense of unity. *: p < 0.05.

Across all time points, higher oxytocin concentrations were positively associated with greater enjoyment (r = 0.21, p = 0.006), stronger unity with players (r = 0.24, p = 0.001), and stronger unity with fans (r = 0.16, p = 0.037; Fig. 3D). Mediation analyses further showed that oxytocin enhanced perceived unity with players via increased enjoyment (indirect effect: z = 2.74, p = 0.006, 95% CI [0.005, 0.023]; Fig. 3E), and similarly enhanced unity with fans through enjoyment (indirect effect: z = 2.61, p = 0.009, 95% CI [0.004, 0.022]; Fig. 3E). These results indicate that baseline-dependent oxytocin dynamics do not merely track social bonding but contribute to it by amplifying enjoyment during spectating, which in turn promotes social bonding among spectators.

### 3.3. Endogenous oxytocin dynamics co-occur with heart rate synchrony

We then investigated whether live sports spectating increases heart rate (HR) synchrony among spectators and whether this synchrony is associated with oxytocin. Mean HR and peak HR followed similar trajectories in the High-OT and Low-OT groups, with both indices showing robust elevations relative to pre-game levels (mean HR: F₂.₅₇,₁₄₆.₂₀ = 84.37, p < 0.0001, ηp² = 0.59; peak HR: F₃.₁₄,₁₈₂.₃₀ = 170.90, p < 0.0001, ηp² = 0.74; Fig. 4B, C). Cross-correlation analyses revealed that HR synchrony, indexed by CCF values, increased significantly during the first and second halves of the game (F₂.₉₇,₁₆₉.₅₀ = 28.19, p < 0.0001, ηp² = 0.33; Fig. 4D). In parallel, the temporal lag between HR time series shortened during and after the game (F₃.₀₃,₁₇₂.₆₀ = 31.39, p < 0.0001, ηp² = 0.36; Fig. 4E), indicating tighter moment-to-moment alignment of cardiac dynamics. Oxytocin concentrations were positively related to HR synchrony (r = 0.15, p = 0.041; Fig. 4F). This association was especially pronounced in the Low-OT group, in which oxytocin levels were significantly associated with both CCF values (r = 0.28, p = 0.009) and time-lag indices (r = −0.26, p = 0.014; Supplementary Fig. 1B, D). These findings suggest that live sports spectating promotes HR synchrony among casual spectators regardless of baseline oxytocin level, and that endogenous oxytocin dynamics, particularly in individuals with lower baseline concentrations, contribute to the processes supporting this synchrony. Importantly, these effects were observed in modestly sized collegiate games, indicating that bonding mechanisms based on HR synchrony generalise beyond the highly fused fan groups typical of professional settings.

**Figure 4.**
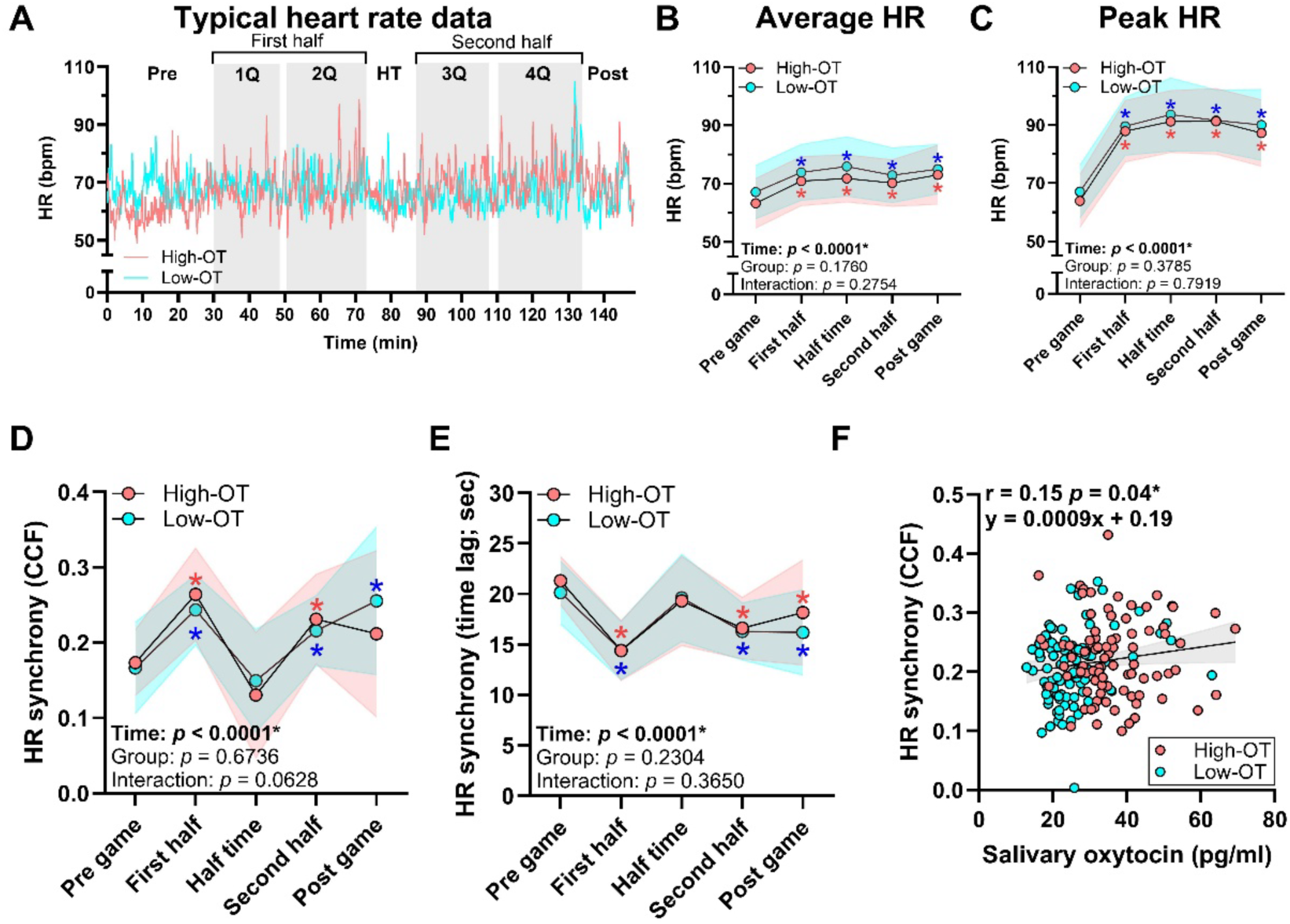
Endogenous oxytocin dynamics co-occur with HR synchrony. **A.** Typical HR data. **B.** Average HR. **C**. Peak HR. **D.** Similarity of HR dynamics (cross-correlation function; CCF). **E.** Time lag. Standard divisions are shown as red (High-OT) bands and blue (Low-OT) on the graph. Results of two-way ANOVA are shown in the lower left of each graph. Time: main effect of watching time. Group: main effect of the group. Interaction: interaction between time and group. *: p < 0.05 vs. pre game for both groups. **F**. Correlation between oxytocin and HR synchrony. The Pearson’s product-rate correlation coefficient and the results of the single regression analysis are shown in the upper-left corner of each graph. The computed regression line and 95% confidence intervals are shown as bands on the graph. *: p < 0.05.

### 3.4. Heart rate synchrony enhances the spectating experience through a sense of unity and loyalty

Finally, we examined how physiological synchrony relates to social bonding and the overall spectating experience, in line with the hypothesis that HR synchrony would support psychological engagement via unity. To characterise the functional role of synchrony, HR data were reclassified into in-game periods (active play) and out-of-game periods (inter-quarter intervals, halftime, and post-game). During in-game periods (Fig. 5A), shorter time-lag values between HR time series were significantly associated with stronger unity toward home team players (r = −0.45, p < 0.001) and higher flow state (r = −0.41, p = 0.001), indicating that tighter moment-to-moment alignment of cardiac dynamics was linked to more affiliative and immersive responses. During out-of-game periods (Fig. 5B), increases in CCF relative to pre-game (ΔCCF) were positively correlated with unity toward home spectators (r = 0.39, p = 0.002) and with loyalty to the team, indexed by intention to revisit (r = 0.27, p = 0.044), suggesting that synchrony outside active play reflects broader social bonding among spectators.

**Figure 5.**
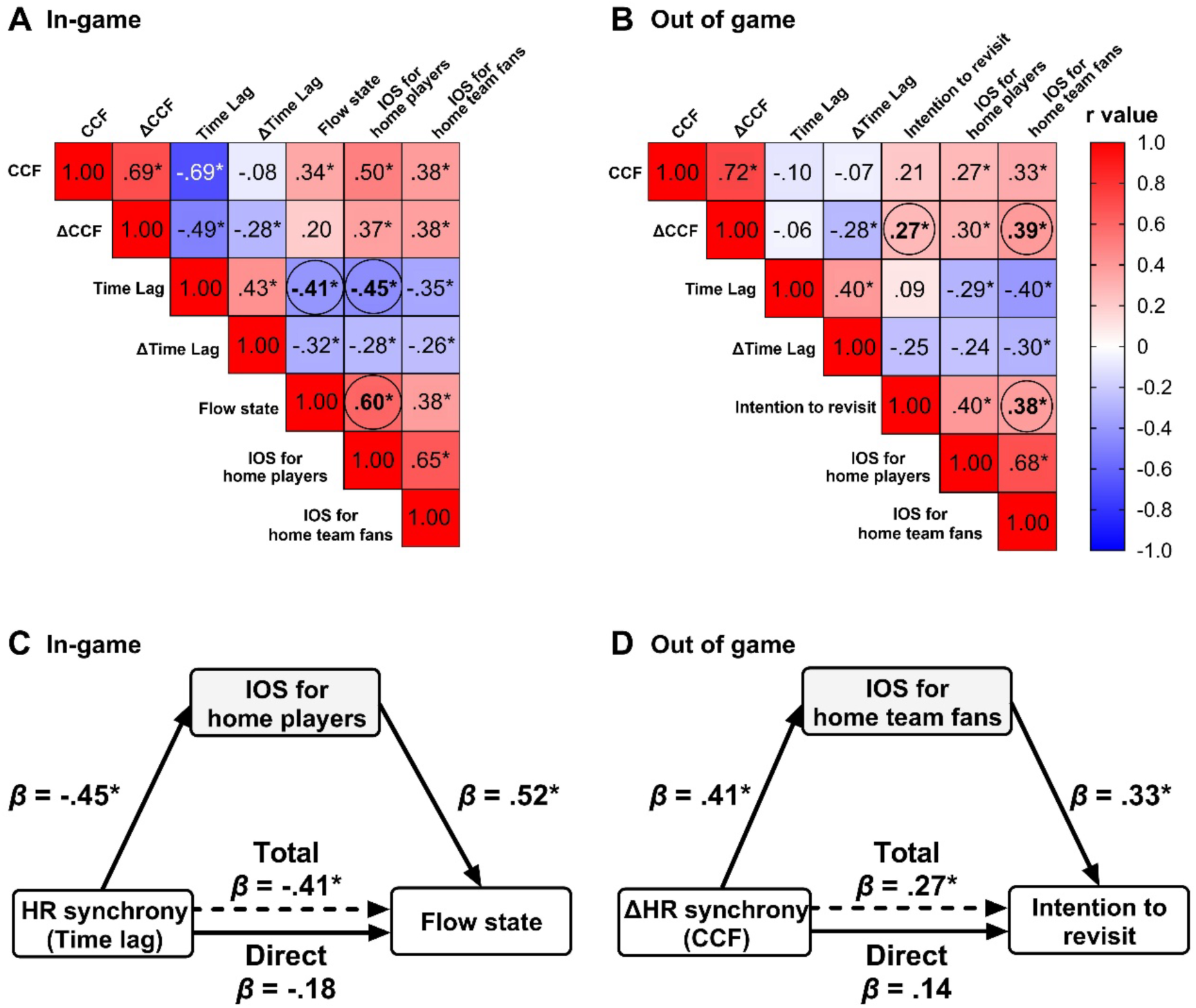
HR synchrony enhances social connection through a sense of unity. **A.** Correlation matrix between measurements related to in-game. **B.** Correlation matrix between measurements related to out of game. **C.** Mediation model of in-game. **D.** Mediation model of out of game. The significance of the correlation analysis is indicated at the top-right of correlation coefficient. * p < 0.05.

To test whether social bonding mediated these relationships, we conducted mediation analyses. In-game, HR synchrony, indexed by shorter time lag, influenced flow state via unity with players as a mediator (z = −3.32, p < 0.001, 95% CI [−0.69, −0.19]; Fig. 5C). Out-of-game, ΔCCF predicted team loyalty via unity with fellow fans (z = 2.01, p = 0.018, 95% CI [0.52, 4.83]; Fig. 5D). These results suggest that physiological synchrony not only reflects but also facilitates the formation of social bonds, which in turn shape meaningful psychological experiences such as immersive flow and loyalty to the team during live spectating, consistent with our hypothesised pathway from oxytocin-linked HR synchrony to social connection.

## 4. Discussion

Live sports spectating in a modest collegiate setting produced two main findings regarding endogenous oxytocin, physiological synchrony, and social bonding among casual spectators. First, oxytocin dynamics during spectating were clearly baseline dependent. Individuals with higher pre-game oxytocin maintained elevated concentrations throughout the game, whereas those with lower pre-game oxytocin showed a marked increase that brought their post-game levels into a similar range (Fig. 2A; Table S1). Across both groups, cortisol decreased and the oxytocin-to-cortisol ratio increased, indicating a shift toward an oxytocin-dominant, low-stress hormonal profile rather than a stress-dominated response to the event (Fig. 2B, C).

Second, endogenous oxytocin and heart rate synchrony jointly supported social connection and experiential outcomes. Higher oxytocin levels were associated with greater enjoyment and stronger unity with players and fellow fans (Fig. 3C–E). Stronger cardiac synchrony was likewise associated with greater unity, higher flow, and stronger loyalty to the team, with distinct in-game and out-of-game relationships (Figs. 4D–F, 5A, B). Mediation analyses further showed that oxytocin and synchrony facilitated these psychological benefits via enhanced feelings of unity, such that enjoyment and unity mediated the links between oxytocin or synchrony and flow or intention to revisit (Figs. 3E, 5C, D). Together, these findings suggest that live sports spectating can act as a naturalistic, non-pharmacological context in which endogenous oxytocin and physiological synchrony are recruited to promote social bonding and positive psychological states in everyday, low-stakes settings.

### 4.2. Baseline-dependent oxytocin dynamics and hormonal profile

The baseline-dependent pattern of oxytocin change observed here complements and extends previous work on endogenous oxytocin responses to affiliative contexts and physical activity. Sports spectating promoted oxytocin dynamics in a manner that depended on baseline levels: individuals with higher pre-game oxytocin maintained elevated concentrations, whereas those with lower pre-game oxytocin showed marked increases that brought their post-game levels into a similar range (Fig. 2A; Table S1). Across both groups, cortisol decreased and the oxytocin-to-cortisol ratio increased (Fig. 2B, C), indicating a shift toward an oxytocin-dominant, low-stress hormonal profile rather than a stress-dominated response to the event.

Previous research has demonstrated that oxytocin release is facilitated by social experiences such as eye contact, shared narratives, and storytelling [36,37]. Our findings suggest that similar mechanisms may be engaged during sports spectating. Elements such as indirect eye contact mediated through players, the embodied performance conveyed by athletes, and the unscripted dramatic nature of the game share key features with storytelling. The collective enjoyment of these elements, as a fundamental human social practice, may serve as a form of “embodied storytelling” that enhances oxytocin dynamics and provides a basis for strengthening social bonds among spectators. In this sense, live sports spectating extends a broader class of everyday affiliative activities, including shared narratives, mutual gaze, and coordinated movement, that are known to upregulate endogenous oxytocin without pharmacological manipulation, offering a behaviourally embedded means of engaging the oxytocinergic system [38–41,37,42].

Mediation analyses further demonstrated that oxytocin fostered unity with players and fellow fans through enhanced enjoyment (Fig. 3E). Taken together, these results align with evidence indicating modest but reliable associations between endogenous oxytocin and social interaction, together with substantial inter-individual heterogeneity and baseline-dependent responsivity [43,44]. The concurrent decrease in cortisol and increase in the oxytocin-to-cortisol ratio also contrast with the cortisol elevations typically observed in competitive athletes and highly invested fans during high-stakes events [45–47], suggesting that modest collegiate spectating can support social engagement within a hormonal milieu that avoids pronounced stress-related endocrine costs.

### 4.3. Neural mechanisms and translational implications of oxytocin dynamics

The observed mediation, whereby oxytocin enhanced unity via enjoyment (Fig. 3E), may be accounted for by the neural mechanism through which oxytocin potentiates activity in the brain’s reward system. Oxytocin receptors are abundantly expressed in regions central to reward processing, including the ventral tegmental area (VTA) and nucleus accumbens (NAc) [48]. Animal studies have shown that oxytocinergic neurons in the hypothalamic paraventricular nucleus project directly to the VTA, where they enhance dopaminergic signalling via oxytocin receptors [49]. Converging evidence in humans indicates that intranasal oxytocin increases activation of the VTA and NAc when participants view a partner’s face or other socially salient cues [50]. These findings support the notion that oxytocin enhances the valuation of social stimuli through reward-related pathways. While sports spectating has long been known to engage the reward system [51], the neural basis of social bonding during collective spectating has remained elusive. Our results suggest that elevated oxytocin during sports spectating may amplify reward system activity as a form of social reward, thereby providing a neural mechanism for fostering unity with players and fellow fans.

Beyond its actions on mesolimbic structures underlying the reward system, oxytocin may also influence higher-order regions of the prefrontal cortex. A prominent candidate is the ventromedial prefrontal cortex (vmPFC). The vmPFC receives projections from the VTA and NAc and is well established as a core component of the reward system [52], yet it is also engaged by others’ successes and socially salient events, implicating it in vicarious reward processing [53]. Recent evidence further indicates that oxytocin enhances vmPFC activity in response to socially salient cues [54,55]. Thus, the facilitation of oxytocin dynamics during sports spectating may strengthen unity with players and fellow fans through modulation of mesolimbic and vmPFC function, linking shared emotional engagement with neuroendocrine and neural mechanisms of social reward.

From a translational perspective, these baseline-dependent and low-stress oxytocin dynamics support the idea that live sports spectating may serve as a naturalistic, non-pharmacological means of recruiting endogenous oxytocin in everyday life. Such an approach could complement pharmacological oxytocin administration by targeting the same neuropeptide system through behavioural rather than exogenous routes, potentially helping to address some of the challenges related to efficacy, dosing, and long-term safety that have emerged in clinical trials of intranasal oxytocin [56,7,5,8,57,11].

### 4.4. Heart rate synchrony as a component of collective experience

This study also demonstrated that live sports spectating promotes heart rate (HR) synchrony among spectators irrespective of baseline oxytocin levels (Fig. 4D, E). Such synchrony is likely driven by the intense collective arousal elicited during spectating. Shared external stimuli, including the tension of the game, the roar of the crowd, and collective reactions, act as powerful forces that align cardiac dynamics across individuals, transcending individual hormonal baselines. Physiological synchrony has similarly been observed in other collective events such as music concerts and religious rituals, where it is regarded as a physiological substrate of shared group experiences [58,59]. Our findings suggest that sports spectating shares this property, functioning like other collective gatherings in fostering synchrony among casual fans.

In line with this literature, HR synchrony increased markedly during the game in our sample, with shorter time lags and higher cross-correlation values indicating tighter temporal alignment of cardiac dynamics among spectators (Fig. 4D, E). Importantly, the present study extends prior work by demonstrating functionally distinct links between synchrony and psychological outcomes in different phases of the event: during in-game periods, higher synchrony, indexed by shorter time lags, was associated specifically with stronger unity toward players and greater flow (Fig. 5A); during out-of-game periods, increases in synchrony relative to pre-game were related to unity with fellow spectators and a stronger intention to revisit (Fig. 5B). These phase-specific associations suggest that synchrony during active play is especially tied to immersive, player-focused engagement, whereas synchrony during breaks and post-game is more closely related to bonding within the spectator community and subsequent loyalty-related motivation.

### 4.5. Coupling between oxytocin, synchrony, and social experience

Another key finding of this study is the observed association between endogenous oxytocin levels and HR synchrony (Fig. 4F). Although both oxytocin release and physiological synchrony are facilitated in collective events and social interactions such as eye contact and touch [58,36,37], their direct coupling has rarely been tested. Experimental work shows that exogenous oxytocin can increase physiological linkage in small groups during cooperative tasks, and research in parent–infant dyads links endogenous oxytocin to interactive synchrony [60,61]. To our knowledge, however, no study has directly linked moment-to-moment fluctuations in endogenous oxytocin to physiological synchrony in a large, naturalistic spectator setting. Evidence that intranasal oxytocin enhances synchrony in laboratory cooperation tasks is therefore mechanistically consistent with our field data [61,62], yet the present association in live sports extends these insights to ecologically valid group contexts.

Mediation analyses further showed that unity statistically mediated the relationships between synchrony and experiential outcomes (Fig. 5C, D). Synchrony enhanced flow via unity with players during active play, and enhanced team loyalty via unity with fellow fans during breaks and post-game. Together with the observation that higher oxytocin levels co-occurred with stronger synchrony and social bonding (Figs. 3D, 4F), these findings support accounts in which endogenous oxytocin facilitates shared attention, emotional convergence, and alignment of autonomic states. In this view, oxytocin-linked synchrony can promote both on-court immersion and off-court social affiliation in spectating contexts. Taken together, our findings suggest a mutually reinforcing relationship between oxytocin and HR synchrony, situating sports spectating alongside other collective gatherings as a naturalistic context that both elicits physiological synchrony and tracks with endogenous oxytocin dynamics.

### 4.6. Broader social and clinical implications

These observations carry translational implications. Oxytocin has long been considered a promising pharmacological candidate for improving social functioning in autism spectrum disorder (ASD) and related conditions, yet large multisite randomised trials have yielded null or modest effects on core social symptoms when intranasal oxytocin is administered over months [8]. In parallel, safety and methods reviews emphasise uncertainties regarding nose-to-brain delivery, individual response profiles, and potential peripheral off-target effects, cautioning against viewing exogenous oxytocin as a simple “social drug” [57]. By contrast, our data show that a brief, enjoyable, and widely accessible group activity, live sports spectating, can engage endogenous oxytocin dynamics and oxytocin-linked HR synchrony in a naturalistic setting, without pharmacological exposure or stress-related hormonal costs (Figs. 2–5). At the population level, sports spectating has been associated with higher well-being and lower loneliness [46,63,64], and ASD research indicates difficulties in spontaneous interpersonal synchrony [65,66], suggesting that synchrony-supportive environments could serve as low-burden adjuncts to social habilitation. While causal clinical benefits remain to be established, the present findings raise the possibility that inclusive spectator contexts might help harness endogenous oxytocin-linked synchrony to scaffold social engagement in populations with social-communication difficulties.

Beyond their immediate relevance to oxytocin biology and physiological synchrony, these findings also inform broader discussions about how everyday collective activities support social well-being. Social psychological models of sport fandom propose that team identification and regular spectatorship enhance well-being by providing opportunities to gain and maintain social connections with other fans [29,30,28]. From this perspective, live sports spectating can be viewed as a form of embodied storytelling, in which the embodied performances of players, the indirect eye contact created by shared attention to the court, and the unscripted drama of the game are collectively experienced and emotionally co-constructed [67,36,37]. Historically, large-scale collective practices ranging from religious rituals and martial displays to seasonal festivals and stadium sports have provided embodied arenas for synchronising bodies and emotions, with modern spectator sports representing a contemporary expression of this long-standing human tendency toward coordinated collective experience [58,68,67,59].

Against the backdrop of Japanese collegiate sports, these findings carry particular significance. In the United States, collegiate sports exert far-reaching social influence, supported by strong athletic performance, substantial financial resources, and extensive entertainment infrastructures. By contrast, Japanese collegiate sports have not yet achieved comparable popularity or institutional support [69]. Nevertheless, our study revealed that even in the relatively local and small-scale context of Japanese university basketball, spectators synchronised, forged social bonds, and developed stronger intentions to revisit (Fig. 5C, D). These results indicate that the social benefits of spectating are not confined to professional sports on grand stages. Rather, they can also emerge from sharing the efforts of familiar individuals, potentially providing a socio-physiological foundation for community building and the promotion of well-being in local societies through sports.

Emerging formats of collective viewing, such as e-sports arenas, large-scale online streaming events, and bio-sharing interfaces that transmit physiological signals between remote spectators, may reproduce or transform the embodied storytelling processes identified here. For example, real-time sharing of bodily signals such as facial expressions and HR in e-sports has been shown to enhance the perceived social presence of others [70]. This suggests that embodiment, a universally available human resource, serves as a foundation for creating social connections. Future work should explore how the sharing of embodiments across time and space can be harnessed to promote collective well-being via endogenous oxytocin dynamics and oxytocin-linked synchrony in increasingly hybrid physical–digital environments.

### 4.7. Limitations and future directions

While our field-based approach ensured ecological validity by capturing real-time physiological dynamics during live sports events, several limitations should be acknowledged. First, the determinants of individual differences in baseline oxytocin levels remain unclear. In this study, participants were categorised based on the median pre-game oxytocin concentration, and we verified that this classification was not attributable to physical characteristics such as body composition, resting HR, or habitual physical activity (Table S1). Nonetheless, it remains uncertain whether the observed variability reflects other stable individual traits or transient influences such as recent social interactions.

Second, oxytocin was measured in saliva, leaving the dynamics of oxytocin within the central nervous system unresolved. Salivary oxytocin tracks peripheral oxytocin to some extent: in mothers, concurrently sampled saliva and plasma concentrations covary across tasks, and a preregistered meta-analysis reports modest central–peripheral coupling that emerges after experimental stimulation rather than at rest [71,72]. In humans, plasma oxytocin has been shown to predict cerebrospinal-fluid oxytocin in a lumbar-puncture study [73], although other reports find no baseline association, underscoring ongoing measurement debate [74].

Accordingly, while salivary assays offer a practical window on the oxytocinergic system in the field, inference to brain dynamics remains indirect. Future studies should combine validated peripheral sampling with neuroimaging and, where feasible, pharmacological challenges to delineate the neural mechanisms by which sports spectating fosters social bonding [12].

Third, our analyses focused on spectator dyads sharing similar attributes, such as seating location and game exposure. Whether HR synchrony also occurs between spectators of different demographics, nationalities, or group affiliations remains unknown. Future research should extend these investigations to diverse spectator pairings and, importantly, to synchrony between spectators and players. Finally, the observational design without a non-spectating control group precludes strong causal inferences about the effects of live sports spectating per se, and unmeasured situational factors, including team performance expectations, social relationships among co-attendees, or pre-existing fan loyalty, may have contributed to both hormonal and psychological responses. Combining experimental or quasi-experimental manipulations with more diverse samples, repeated events, and longitudinal follow-up will be essential for testing whether oxytocin-linked synchrony during live spectating prospectively predicts sustained changes in social connection, mental health, and engagement in community activities.

## 5. Conclusion

This study showed that a single session of live sports spectating in a real-world collegiate setting can recruit endogenous oxytocin and interpersonal heart rate synchrony in ways that support social bonding among casual fans. Spectators with low baseline oxytocin exhibited clear increases in oxytocin and a shift toward an oxytocin-dominant, low-stress hormonal profile, while higher oxytocin levels and stronger heart rate synchrony across participants were associated with greater enjoyment, stronger unity with players and fellow fans, and higher flow and intention to revisit. These findings suggest that live sports spectating may serve as a simple, scalable non-pharmacological approach to engaging oxytocin-related systems and interpersonal synchrony to promote social connection in everyday life. Such contexts may in turn encourage sustained community engagement and contribute to more active, healthier, and more socially connected lifestyles.

## Acknowledgements

We would like to thank all staff at the Bureau of Physical Education and Sports, University of Tsukuba, for their dedicated efforts in organizing the home series event “Tsukuba Live!” This research was supported by a part of ‘Sports in Life’ project by the Japan Sports Agency to K.K. and T.M.; a Special Research Grant from the TOYOTA Foundation to T.M. (D20-ST-0034); a Fusion Oriented Research for disruptive Science and Technology (FOREST) by Japan Science and Technology Agency (JST) to T.M. (JPMJFR205M).

## Data availability

The data supporting the findings of this study are available from the corresponding author upon reasonable request.

## Author contributions

T.M., T.Y., and K.K. conceived and designed the study. T.M., T.Y., S.T., H.M. S.D., K.K., S.Y., and H.T. recruited participants. T.M., T.Y., D.F., S.T., H.M., S.D., and K.K. collected the data. T.M., T.Y., D.F., S.T., H.M., and S.D. conducted the data analysis and interpreted the data. T.M. and T.Y. drafted the manuscript. T.M., T.Y., D.F., S.T., S.D., K.K., H.T. edited and revised the manuscript. All authors have read and approved the final version of the manuscript.

## Declaration of generative AI in scientific writing

During the preparation of this work the authors used ChatGPT for only English editing. After using this service, the authors reviewed and edited the content as needed and took full responsibility for the content of the published article.

---

**Table S1.** Participants’ characteristics.

|  | **High-OT**<br>**(n = 30)** | **Low-OT**<br>**(n = 30)** | ***p*** | ***d*** |
| --- | --- | --- | --- | --- |
| Median oxytocin (pg/ml) | 27.5 |  |  |  |
| Baseline oxytocin levels (pg/ml) | 31.9 ± 9.2<sup>#</sup> | 21.6 ± 3.7 | <0.01 | 2.46 |
| Age (years) | 22.2 ± 8.6 | 22.9 ± 9.0 | 0.77 | 0.07 |
| Gender (n, Male/Female) | 20 / 10 | 19 / 11 |  |  |
| Height (cm) | 172.3 ± 9.8 | 168.6 ± 7.4 | 0.11 | 0.42 |
| Body weight (kg) | 68.6 ± 15.8 | 65.3 ± 8.8 | 0.34 | 0.25 |
| BMI (kg/m<sup>2</sup>) | 23.3 ± 1.2 | 22.9 ± 3.3 | 0.71 | 0.11 |
| Resting HR (bpm) | 63.2 ± 8.5 | 67.1 ± 9.4 | 0.10 | 0.43 |
| Vigorous-intensity activity<br>(min/day) | 87.3 ± 82.6 | 80.7 ± 80.8 | 0.76 | 0.08 |
| Moderate-intensity activity<br>(min/day) | 16.8 ± 18.9 | 20.4 ± 39.7 | 0.66 | 0.11 |
| Walking time (min/day) | 24.1 ± 29.6 | 28.1 ± 40.3 | 0.67 | 0.11 |
| Exercise time (min/day) | 70.0 ± 67.7 | 70.5 ± 83.3 | 0.97 | <0.01 |
| Sitting time (min/day) | 348.6 ± 226.6 | 284.5 ± 160.9 | 0.23 | 0.32 |
| Cheering for home team (score) | 6.6 ± 0.5 | 6.2 ± 0.9 | 0.10 | 0.43 |

Values are presented as mean ± SD. #: $p < 0.05$ vs. Low-OT.

**Figure S1.**
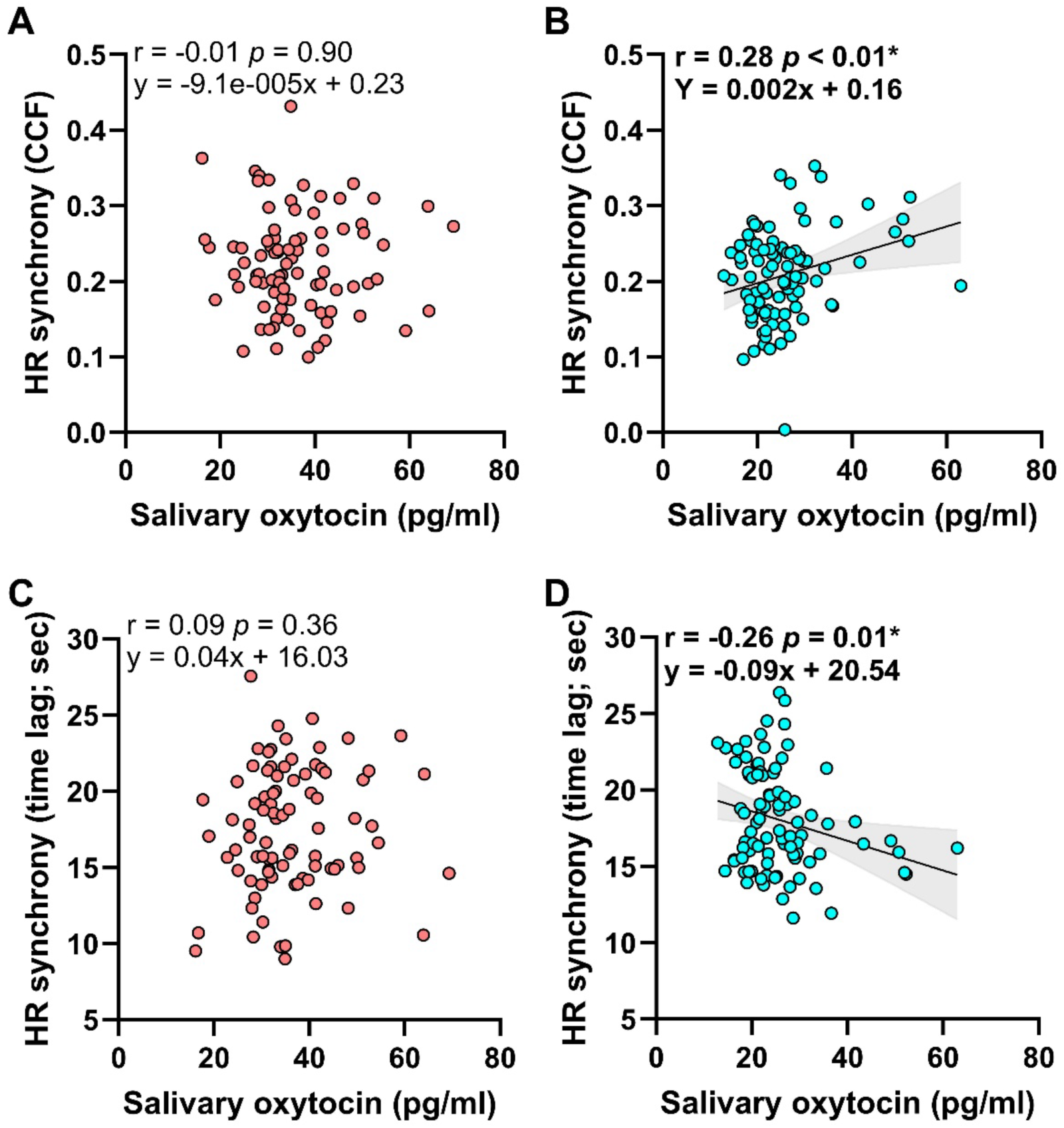
Stronger relationship between HR synchrony and oxytocin in the Low-OT group. **A, B.** Correlation between oxytocin and HR synchrony (CCF) in High-OT group (A), and Low-OT group (B). **C, D**. Correlation between oxytocin and HR synchrony (time lag) in High-OT group (C), and Low-OT group (D). The Pearson’s product-rate correlation coefficient and the results of the single regression analysis are shown in the upper-left corner of each graph. The computed regression line and 95% confidence intervals are shown as bands on the graph. *: p < 0.05.

**Figure S2.**
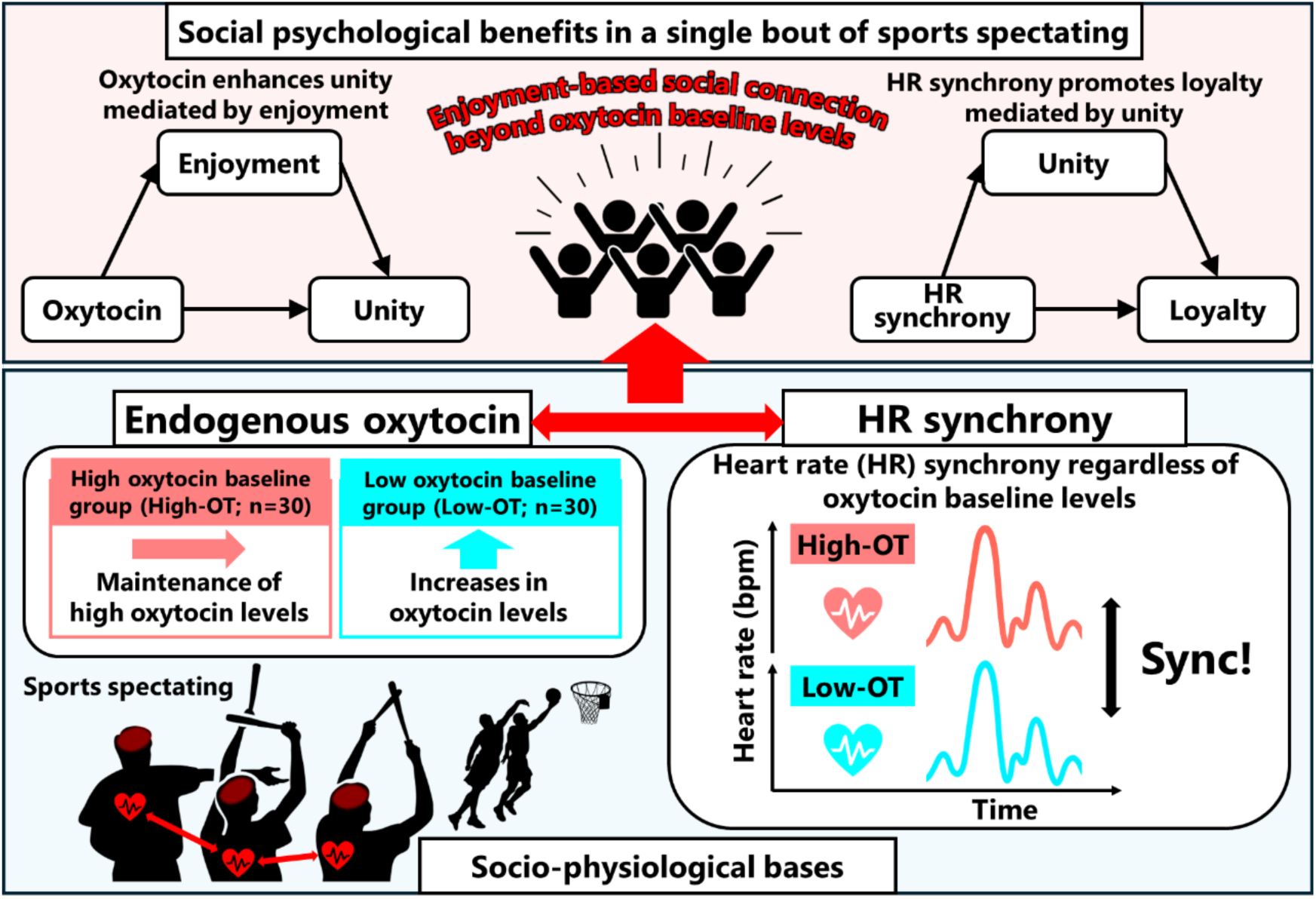
Graphical model of how sports spectating may foster enjoyment-based social connection via socio-physiological responses. Sports spectating promotes baseline-dependent oxytocin dynamics: individuals with high baseline oxytocin maintain elevated concentrations, whereas those with low baseline levels exhibit increases during spectating. These oxytocin changes, in turn, facilitate heart rate synchrony regardless of baseline levels, and the two processes may reciprocally reinforce each other. Such socio-physiological bases ultimately foster enjoyment-based social connection among spectators.

